# COATi: statistical pairwise alignment of protein-coding sequences

**DOI:** 10.1101/2023.05.22.541791

**Authors:** Juan J. García Mesa, Ziqi Zhu, Reed A. Cartwright

## Abstract

Sequence alignment is an essential method in bioinformatics and the basis of many analyses, including phylogenetic inference, ancestral sequence reconstruction, and gene annotation. Sequence artifacts and errors made during alignment reconstruction can impact downstream analyses leading to erroneous conclusions in comparative and functional genomic studies. For example, abiological frameshifts and early stop codons are common artifacts found in protein coding sequences that have been annotated in reference genomes. While such errors are eventually fixed in the reference genomes of model organisms, many genomes used by researchers contain these artifacts, and researchers often discard large amounts of data in comparative genomic studies to prevent artifacts from impacting results. To address this need, we present COATi, a statistical, codon-aware pairwise aligner that supports complex insertion-deletion models and can handle artifacts present in genomic data. COATi allows users to reduce the amount of discarded data while generating more accurate sequence alignments.

## Introduction

Sequence alignment is a fundamental task in bioinformatics and a cornerstone step in comparative and functional genomic analysis (Rosenberg, 2009). While sophisticated advancements have been made, the challenge of alignment inference has not been fully solved (Morrison, 2015). The alignment of protein-coding DNA sequences is one such challenge, and a common approach to this problem is to perform alignment inference in amino-acid space (e.g. Bininda-Emonds, 2005; Abascal et al., 2010). While this approach is an improvement over DNA models, it discards information, underperforms compared to alignment at the codon level, and fails in the presence of artifacts, such as frameshifts and early stop codons. While some aligners can utilize codon substitution models, they are often not robust against coding-sequence artifacts.

Within protein-coding sequences, indels may occur in between any pair of adjacent nucleotides, and therefore, gaps in alignments of natural sequences may occur both between and within codons (Fig. 1). Gaps that occur after the first position or second position of a codon are known as phase-1 and phase-2 gaps, respectively. Gaps that occur between codons are known as either phase-3 gaps (this study) or phase-0 gaps (e.g. Taylor et al., 2004). While all three phases of gaps occur in natural sequences, alignments performed in amino-acid or codon space force all gaps to be phase-3 gaps. Because only about 42% of indels are phase-3 (Taylor et al., 2004; Zhu, 2022), this mismatch between aligner assumptions and biology can produce sub-optimal alignments and inflated estimates of sequence divergence (Fig. 1).

**Figure 1:**
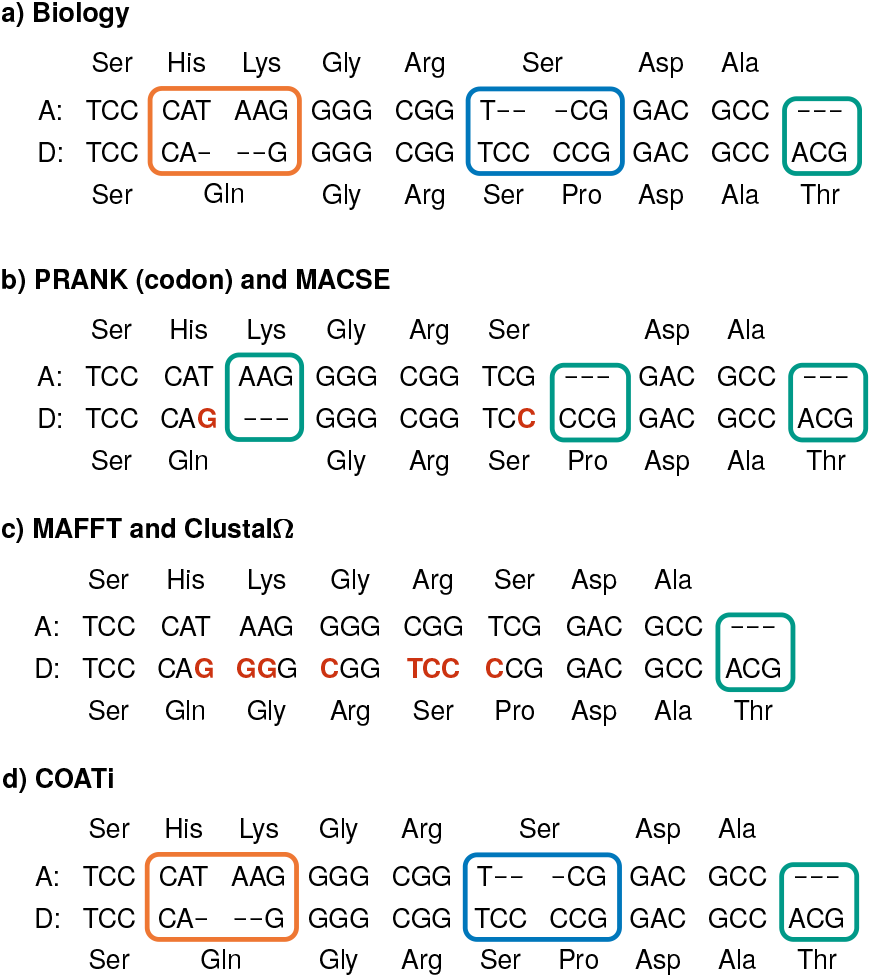
Standard algorithms produce suboptimal alignments. (a) shows the true alignment of an ancestor sequence (A) and a descendant sequence (D). (b–d) are the results of different aligners. Nucleotide mismatches are highlighted in red.—Notably, COATi is the only aligner able to retrieve the biological alignment in this example.—Indels in protein-coding sequences can be classified as having one of three difference phases and being one of two different types. Phases refer the the location of the gap with respect to the reading frame, while types refer to the consequence of the indel. Phase-1, phase-2, and phase-3 indels are shown in blue, orange, and green respectively. Additionally, the orange indel is type-II (an amino-acid indel plus an amino-acid change) while the blue indel is type-I (an amino-acid indel only). The difference between an in-frame and a frameshift indel is not displayed.

Bioinformatic pipelines need to be robust to variation in quality across genomic datasets because uncorrected errors in the alignment stage can lead to erroneous results in comparative and functional genomic studies (Schneider et al., 2009; Fletcher and Yang, 2010; Hubisz et al., 2011). While genomes for model organisms often get refined over many iterations and contain meticulously curated protein-coding sequences, genomes for non-model organisms might only receive partial curation and typically have lower quality sequences and annotations. These genomes often lack the amount of sequencing data needed to fix artifacts, including missing exons, erroneous mutations, and indels (Jackman et al., 2018). When comparative and functional genomics studies include data from non-model organisms, care must be taken to identify and manage such artifacts; however, current alignment methods are ill-equipped to handle common artifacts in genomic data, requiring costly curation practices that discard significant amounts of information.

To address current limitations of alignment software to accurately align protein-coding sequences, we present COATi, short for COdon-aware Alignment Transducer, a pairwise statistical aligner that incorporates evolutionary models for protein-coding sequences and is robust to artifacts present in modern genomic data sets.

## Methods

### Statistical Alignment via Finite State Tranducers

In statistical alignment, sequence alignments are scored based on a stochastic model, typically derived from molecular evolutionary processes (Lunter et al., 2005). An advantage of statistical alignment is that its parameters are derived from biological processes, allowing them to be estimated directly from data or extracted from previous studies. While approaches vary, a statistical aligner for a pair of sequences, *X* and *Y*, typically finds an alignment, *Aln*, that maximizes the joint probability *P* (*Aln, X, Y*) or samples alignments from the posterior *P* (*Aln*|*X, Y*). This is typically performed using pairwise hidden Markov models (pair-HMMs; Bradley and Holmes, 2007). Pair-HMMs are computational machines with two output tapes. Each tape represents one sequence, and a path through the pair-HMM represents an alignment of the two sequences. Conceptually, pair-HMMs generate two sequences from an unknown ancestor and can calculate the joint probability *P* (*Aln, X, Y*) (Yoon, 2009).

While the use of pair-HMMs is ubiquitous in bioinformatics, they are limited to modeling the evolution of two related sequences from an unknown ancestor. As an alternative, finite-state transducers (FSTs, Fig. 2) allow researchers to model the evolution of a descendant sequence from an ancestral sequence. FSTs are computational machines with one input tape and one output tape and provide similar benefits to pair-HMMs, while being more suitable for evolutionary models (Bradley and Holmes, 2007). FSTs consume symbols from an input tape and emit symbols to an output tape based on the symbols consumed and the structure of the FST. Conceptually, FSTs generate a descendant sequence, *D*, from a known ancestor, *A*, and can calculate the conditional probability *P* (*Aln, D*|*A*).

**Figure 2:**
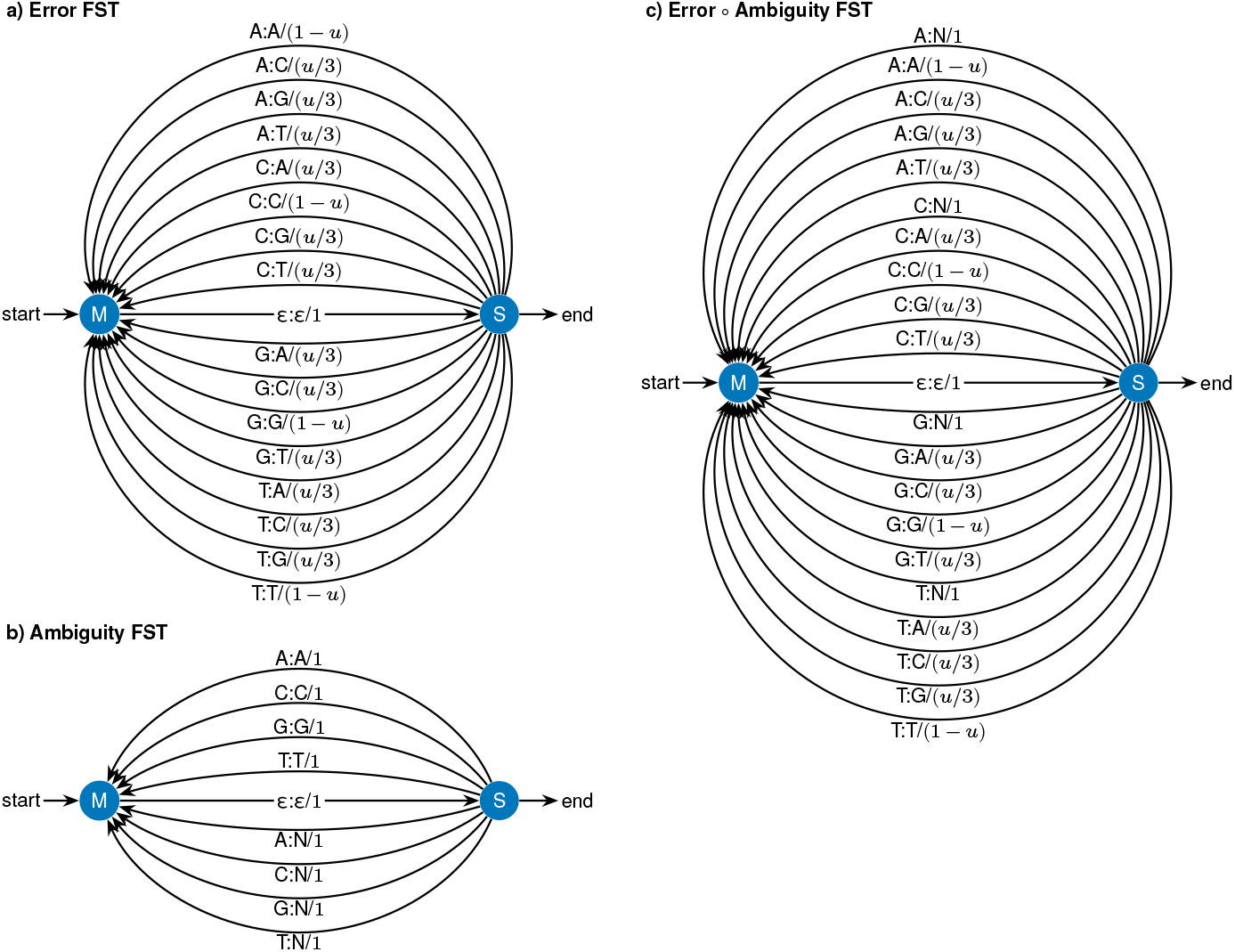
Finite state transducers (FSTs) model the generation of an output sequence based on an input sequence. (a) A graph of a probabilistic FST (Cotterell et al., 2014) for base-calling errors using a Mealy-machine architecture, where parameter *u* is the error rate. This graph contains two states (S and M) connected by arcs, with labels “input symbols : output symbols / weight”. Arcs consume symbols from the input sequence and emit symbols to the output sequence. Weights describe the probability that an arc is taken given the input symbols. Epsilon (ε) is a special symbol denoting that no symbols were either consumed or emitted. (b) An FST for matching sequences against ambiguous nucleotides (N). (c) An FST that results from the composition (∘ operation) of the Error FST with the Ambiguity FST.

There are well-established algorithms for combining FSTs in different ways allowing the design of complex models by combining simpler FSTs, including concatenation, composition, intersection, union, and reversal (Bradley and Holmes, 2007; Silvestre-Ryan et al., 2021). Specifically, composition is an algorithm to combine two FSTs by sending the output of one FST into the input of another, creating a new, more complex transducer (Mohri et al., 2005). Figure 2 illustrates how FSTs modeling sequencing errors (Fig. 2a) and ambiguity (Fig. 2b) can be combined via composition to produce an FST that does both (Fig. 2c). Conceptually, composition creates an FST that generates a descendant sequence from a known ancestor via an unknown intermediate, *J*, and can calculate the conditional probability *P* (*D*|*A*) = ∑ _*J*_ *P* (*D*|*J*)*P* (*J*|*A*).

### The COATi FST

COATi aligns pairs of sequences using a statistical alignment model, which is implemented as a finite-state transducer derived from the composition of multiple FSTs, each representing a specific biological or technical process: (1) the codon substitution FST, (2) the indel FST, (3) the error FST, and (4) the ambiguity FST (Figs. 2–3; c.f. Holmes and Bruno, 2001). We call this transducer the COATi FST. Codon substitution models are uncommon in sequence aligners, despite their extensive use in phylogenetics. COATi implements both the Muse and Gaut (1994) codon model (codon-triplet-mg) and the Empirical Codon Model (codon-triplet-ecm; Kosiol et al., 2007). It also lets the user provide a codon substitution matrix. A key innovation of COATi is that it combines a codon substitution model with a nucleotide-based indel model, allowing gaps to occur both between and within codons (c.f. Ranwez et al., 2011, 2018). This also allows the aligner to be robust against sequencing artifacts that produce sequences with disrupted reading frames.

**Figure 3:**
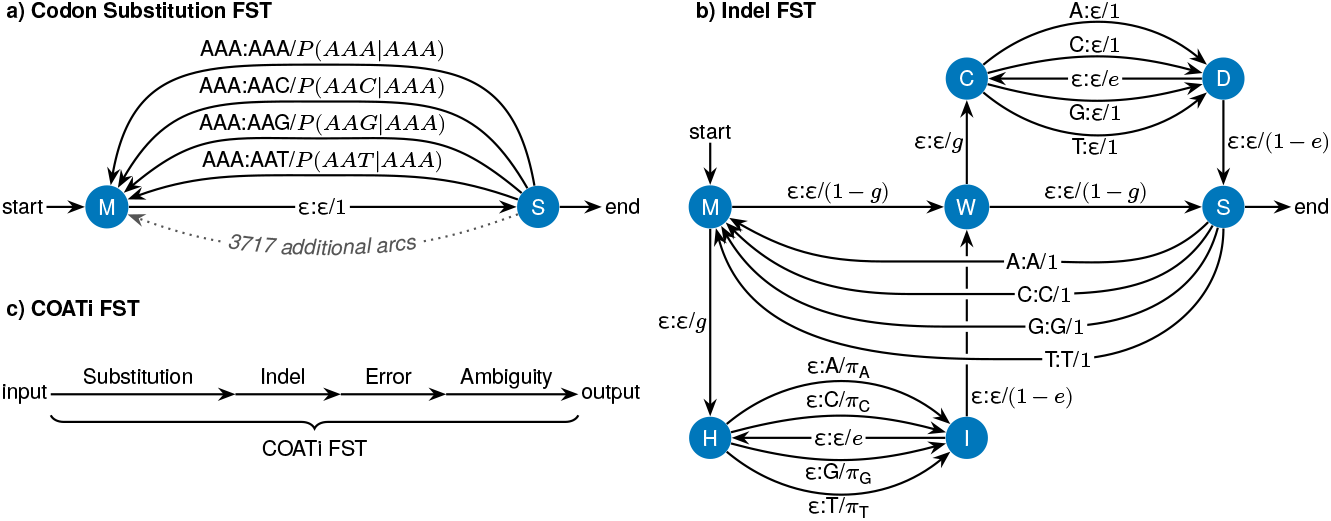
The COATi FST is built from simpler FSTs via composition. (a) The substitution FST encodes a 61 × 61 codon substitution model with 3721 arcs from S to M. These arcs consume three nucleotides from the input tape and emit three nucleotides to the output tape. The weight of each arc is a conditional probability derived from a codon substitution model. See Fig. 2 for more details about reading this graph. (b) The indel FST allows for insertions (H to I) and deletions (C to D). Here *g* is the gap-opening parameter and *e* is the gap-extension parameter. Insertion arcs are weighted according to the codon model’s stationary distribution of nucleotides, and deletion arcs have a weight of 1. This FST is structured such that if insertions and deletions are contiguous, insertions will precede deletions. (c) The COATi FST is derived via composition from the codon substitution, indel, error, and ambiguity FSTs.

COATi FST is not a true probabilistic FST and cannot be used as-is to simulate output sequences based on an input sequence. This is because it is missing a parameter to control how often Ns are added to the output sequence. This is a design feature to allow COATi FST to properly weight ambiguous nucleotides as representing any other symbol. In addition, COATi’s indel FST (Fig. 3b) implicitly assumes that match states occur both immediately before and immediately after the alignment. This allows COATi FST to assign the same weight to gaps at the beginning and end of an alignment but also introduces a normalization constant (not shown) to reflect that mass is lost by not allowing the indel FST to terminate from node C. While this normalization constant is not needed to find the most likely alignment or to sample alignments, it can be calculated using a Markov model that considers only the transitions between states M and D in the indel FST (Fig. 3b) and ignoring any states in between:

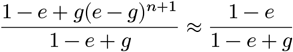

where *g* is the gap opening parameter, *e* is the gap-extension parameter, and *n* is the length of the input sequence. The exponent has a +1 due to the presence of an “immortal match” immediately after the input sequence.

Since codon substitution is the first process in COATi’s model, the input sequence to the COATi FST must be compatible with a codon-substitution model and be a multiple of three nucleotides and not contain any ambiguous symbols or stop codons. The phases, reading frames, and amino-acid contexts of alignment columns is determined by the input sequence, and better alignments will be generated if the input is of high quality and free of artifacts. Depending on context, we may refer to the input sequence as the “ancestral” or “reference” sequence. In contrast, the output sequence must be compatible with the ambiguity FST, can be of any length, and can contain any nucleotides or “N”. This allows COATi to align lowerquality sequences that may contain artifacts against a high-quality reference sequence. We may refer to the output sequence as the “descendant” or “non-reference” sequence. The choice of which sequence is the input sequence and which sequence is the output sequence is left up to the user.

In order to use the COATi FST to align an output sequence against an input sequence, we first convert each sequence into an acceptor, represented as a linear transducer where the input and output symbols of each transition are identical and each transition represents one nucleotide of a sequence (Allauzen et al., 2007). By composing the input and output acceptors with the COATi FST, we generate a transducer of all possible alignments of the two sequences. Any path through this FST represents a pairwise alignment, while the shortest path (by weight) corresponds to the best alignment. If more that one optimal alignment exists, ties are broken according to the implementation of the shortest-path algorithm. All FST operations in COATi, including model development, composition, search for the shortest path, and other optimization algorithms, are performed using the C++ openFST library (Allauzen et al., 2007). An example of an FST-based alignment can be found in Supplementary Materials Figure S1.

### The Marginal Model

The COATi FST has a large state space to keep track of codon substitution rates when codons can be interspersed with indel events. This additional state space increases the computational complexity of the alignment algorithm. To reduce the runtime complexity of COATi, we have also developed an approximation of the COATi FST that can be implemented with standard dynamic programming techniques. This approximation uses a marginal substitution model where the output nucleotides are independent of one another and only depend on the input codon and position. This produces a (61 × 3) × 4 substitution model and eliminates the need to track dependencies between output nucleotides.

A marginal substitution model is calculated from a standard substitution model by calculating the marginal probabilities that each ancestral codon produces specific descendant nucleotides at each reading frame position. Specifically, let

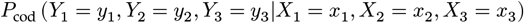

represent transition probabilities from a codon model, and

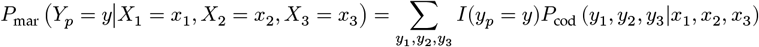

represent the marginal transition probabilities, where *p* ∈ {1, 2, 3} is the position of the descendant nucleotide relative to the ancestral reading frame and *I*() is an indicator function. COATi contains marginal models for both Muse and Gaut (1994) and the Empirical Codon Model (Kosiol et al., 2007), resulting in the marginal models codon-marginal-mg and codonmarginal-ecm. These models emphasize the position in a codon where the substitution occurs, help restrict the effects of low-quality data in the descendant sequence, and allow more than one substitution per codon. In combination with the indel model, alignment using the marginal model is implemented using dynamic programming.

### Empirical Dataset and Alignments

Humans and gorillas are two closely related species with very different levels of genome curation. The human reference genome has been revised dozens of times and is currently on version GRCh38.p14, while the gorilla reference genome has only been revised a handful of times and is currently on version gorGor4 (cf. ENSEMBL database v110; Hubbard et al., 2002). Additionally significant levels of investment have been made to correctly identify and annotate human genes, while gorilla annotations have received limited support in comparison. Together, these reference genomes provide a good opportunity to compare COATi against other aligners as they offer one genome that is high-quality (human) and sister genome that is lower-quality (gorilla).

We used ENSEMBL database v110 (Hubbard et al., 2002) to create an empirical dataset of protein-coding sequences for both human genes and their gorilla orthologs. We first selected human protein-coding genes that belonged to the Consensus Coding Sequence Set, that were located on an autosomal chromosome, and that had a one-to-one gorilla ortholog. We selected the canonical isoform for both species and removed any pair in which the total nucleotide length was larger larger than 6,000 nucleotides. This resulted in 14,127 sequence pairs and corresponding FASTA files containing CDS sequences. Due to the way that canonical isoforms are identified, there is no guarantee that the isoforms are orthologous even though the genes are. Therefore, a subset of the sequence pairs in this dataset contain human and gorilla sequences with different exon compositions. We have made no attempt to correct these sequence pairs because in our experience genome-wide studies rarely control for such artifacts.

### Alignment Methods

In order to compare COATi against other aligners, we evaluated five different alignment models: (1) COATi’s FST model (i.e. codon-triplet-mg), (2) the amino-acid model of ClustalΩ (Sievers et al., 2011) via translation and reverse translation, (3) the amino-acid aware nucleotide model of MACSE (Ranwez et al., 2018), (4) the DNA model of MAFFT (Katoh and Standley, 2013), and (5) the codon model of PRANK (Löytynoja, 2014). COATi is not a symmetric pairwise aligner, as reference sequences are more constrained than non-reference sequences. In order to evaluate the importance of the choice for reference sequence, we also aligned sequences pairs using (6) COATi with gorilla sequences as the reference (i.e. COATi-rev). Together, these six methods allow us to evaluate both different alignment strategies and different software implementations. See Supplemental Materials for additional details, including results for COATi’s marginal and ECM models.

We aligned our dataset of human-gorilla orthologs using all six alignment methods and calculated multiple biological and technical statistics on each alignment in order to compare how different alignment methods influence biological conclusions. Additionally, we checked alignments to ensure that our pipeline did not introduce any artifacts into the results, including unexpected characters, empty columns, and aligned sequences of different lengths. We also generated checksums of our sequences with gaps removed to ensure that they were not modified during alignment.

### Evolutionary Distances and Gaps

Alignment inference impacts the estimation of evolutionary distances. To quantify the impact of aligners on the estimation of evolutionary distances, we calculated Kimura’s 2-parameter distance (K2P; Kimura, 1980) for each sequence-pair and aligner combination. K2P corrects for multiple mutations at a site and takes into account differences in transition and transversion rates. It also assumes equal nucleotide frequencies and no variation in the rate of substitution across sites. While Kimura’s 2-parameter distance is more suitable for non-coding sequences, it is straight forward to calculate and provides a quantitative measure of the evolutionary divergence between sequences. We used the R software package ape (Paradis and Schliep, 2019) to calculate K2P distances. Since aligners influence evolutionary distances based on their tendencies to insert gaps into sequences, we also quantified the phases and lengths of gaps introduced by each method as well as the fraction of nucleotides that are aligned against a match, mismatch, or gap.

### Selection

Alignment inference also impacts the identification of genes that have experienced positive and negative selection. To evaluate the impact of aligners on selection identification, we estimated *K*_*S*_ and *K*_*A*_ statistics (also known as *d*_*s*_ and *d*_*n*_) for each estimated alignment. Briefly, *K*_*S*_ is the number of substitutions per synonymous site and *K*_*A*_ is the number of substitutions per non-synonymous site between two protein-coding sequences. We used the method developed by Li (1993) and independently derived by Pamilo and Bianchi (1993) to estimate these statistics as implemented in the R package seqinr (Charif and Lobry, 2007).

Briefly, this method takes two aligned sequences and classifies the sites in each sequence as non-degenerate, two-fold degenerate, or four-fold degenerate based on the standard genetic code. A site is non-degenerate if 0/3 possible nucleotide changes to that site are synonymous, two-fold degenerate if 1/3 of the possible changes are synonymous, and fourfold degenerate if 3/3 possible changes are synonymous. (The rare three-fold degenerate sites are treated as two-fold degenerate in this method.) First the method calculates the average numbers of sites of each type in the sequence pair (*L*_0_, *L*_2_, and *L*_4_). Next, following Li et al. (1985), it uses Kimura’s two-parameter model (Kimura, 1980) and the alignment to estimate the numbers of transitions (*A*_*i*_) and transversions (*B*_*i*_) that occurred per each *i*-th type. Using these statistics, Li (1993) estimated *K*_*S*_ and *K*_*A*_ as

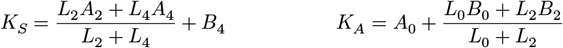

We considered any alignment to be showing evidence of positive selection if *K*_*A*_/*K*_*S*_ > 1 and negative selection if *K*_*A*_/*K*_*S*_ < 1. We estimated F_1_ scores for both positive and negative selection by comparing estimated alignments to benchmark alignments. F_1_ is the harmonic mean of precision and recall:

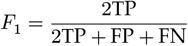

where TP is the number of alignments that correctly predicted positive (or negative) selection, FP is the number of alignments that incorrectly predicted positive (or negative) selection, and FN is the number of alignments that incorrectly predicted the absence of positive (or negative) selection. F_1_ allows us to measure how well aligners produce alignments that correctly identify the presence and absence of positive selection or negative selection.

### Semi-Empirical Benchmark Alignments

In order to compare COATi against other aligners on realistic datasets with known alignments, we developed a procedure to introduce realistic gap patterns into human-gorilla orthologous gene pairs that did not previous contain indels. We separated our 14,127 empirical sequence pairs into two sets: (1) 8261 sequence pairs that did not contain any gaps when aligned by COATi FST, ClustalΩ, MACSE, MAFFT, and PRANK, and (2) 5821 sequence pairs for which at least one of these five aligners added gaps. (45 pairs that could not be aligned by at least one aligner were dropped.) We identified gap patterns from the sequence pairs that were aligned with gaps and randomly introduced them into the ungapped sequence pairs. We used an equal number of randomly sampled gap patterns from each aligner to produce a benchmark set of known alignments. The phases of gaps were preserved as were the spacings of any clusters of gaps. Segments of matches that were 96 nucleotides or longer were allowed to change length to accommodate the length of the ungapped sequence pair. (This criteria was recursively lowered if gap pattern did not fit the ungapped sequence pair.)

For example, consider the gapped alignment represented by the CIGAR string “170M 3D 10M 6I 102M” applied to an ungapped alignment of human and gorilla sequences that are 300 nucleotides long. First, we modified the CIGAR string to insert flexible lengths for any match segment that is 96 or more nucleotides long while preserving phase. This results in a new CIGAR string of “98M *M 3D 10M 6I 96M *M”, where * represents the locations that have flexible length. Considering only matches and deletions, this CIGAR string has 207 fixed nucleotides, leaving 93 nucleotides in the human sequence to be allocated to the flexible locations. Since there are two locations of flexible length, we drew one random break point uniformly while maintaining phase, producing the final CIGAR string of “98M 12M 3D 10M 6I 96M 81M”, which was used to add gaps into the target, ungapped sequence pair. To apply deletions, we removed the corresponding nucleotides from the gorilla sequence, and to apply insertions, we added nucleotides to the gorilla sequence by sampling codons from their stationary frequency while respecting gap phases. The human sequence was left unchanged. We generated a benchmark of 8261 sequence pairs with known alignments using this strategy.

### Measuring Aligner Accuracy

We aligned our benchmark alignments using our suite of aligners and quantified the similarity of each alignment that was produced to its respective benchmark. As above, we measured Kimura’s 2-parameter distance (Kimura, 1980) and *K*_*A*_/*K*_*S*_ (Li, 1993) for the benchmark and estimated alignments. Additionally, we quantified the error of estimated alignments using the *d*_*seq*_ metric (Blackburne and Whelan, 2011).

Intuitively, *d*_*seq*_ ranges between zero and one and can be interpreted as the fraction of nucleotides in the sequence pair that are aligned differently between estimated and benchmark alignments. This metric summarizes each alignment by building homology sets for each nucleotide in the sequence pair. Briefly, if an alignment column contains position *i* from the first sequence and position *j* from the second sequence, then {*j*} is the homology set for position *i* in the first sequence and {*i*} is the homology set for position *j* in the second sequence. If a position is aligned against a gap, then its homology set is simply {gap}. This treats all gaps equally as the location of the gap is not recorded. Finally, *d*_*seq*_ is calculated between an estimated and a benchmark alignment as the average, normalized Hamming distance between the homology sets for each nucleotide in the sequence pair (Blackburne and Whelan, 2011).

Using *d*_*seq*_ we quantified the number of perfect, imperfect, and best alignments each aligner produced. We define perfect alignments as alignments with a distance of zero to the benchmark alignment (*d*_*seq*_ = 0) or any alignment that is evolutionary equivalent to an alignment with a distance of zero. We included equivalency in our definition of perfect alignments to prevent the manner by which aligners break ties from influencing our results. Evolutionary equivalence was determined by scoring alignments using COATi’s marginal model. Any alignment that had a score which matched the score of its benchmark alignment was considered perfect even if its distance to the benchmark was greater than zero. Imperfect alignments are defined as alignments that are not perfect when another method successfully produces a perfect alignment for the same pair of sequences. Best alignments are the alignments that the lowest distance *d*_*seq*_ to the true alignment, including ties and equivalent alignments. Taken together, these three statistics not only allow a direct comparison of aligners but also expose instances where all aligners fall short of achieving a perfect result.

## Results

### Empirical Data and Alignments

We generated 14,127 sequence pairs containing the coding sequences of a human gene and the orthologous gorilla gene. 22 human sequences contained early stop codons. These were not artifacts, but rather UGA codons which encoded for selenocysteine. 1 human sequence had a length that was not a multiple of 3, and no human sequences contained ambiguous nucleotides. Conversely, no gorilla sequences contained early stop codons (i.e. proteins containing selenocysteines were improperly annotated). 20 gorilla sequences had lengths that were not multiples of 3, and 173 sequences contained ambiguous nucleotides. As expected, COATi was not able to align 23 sequence pairs and COATi-rev was not able to align 193 sequence pairs. PRANK was not able to align 24 sequence pairs. MAFFT, MACSE, and ClustalΩ were able to align all sequence pairs, although the latter received some help from its wrapper script. Additionally, MAFFT produced alignments with columns that contained only gaps, which were removed by its wrapper script. A total of 10,296 sequence pairs were aligned equivalently by all five methods, including 8,261 sequence pairs that did not contain any gaps, 664 that had identical alignments, and another 1,371 that had identical scores.

Compared to other aligners, COATi inferred more indels and aligned more nucleotides against a gap while producing fewer mismatches (Tab. 1). Additionally, COATi inferred that 60% indels occurred in phase 1 or phase 2 in our coding sequences. MACSE’s amino-acid aware nucleotide model inferred 7%, while MAFFT’s non-coding DNA model inferred 48%. PRANK’s codon and ClustalΩ’s amino-acid models were constrained by their assumptions and inferred no phase 1 or phase 2 gaps. Note that ClustalΩ’s gaps with phases of 1 and 2 were created by our wrapper script handling DNA sequences with lengths that are not multiples of 3.

**Table 1:**
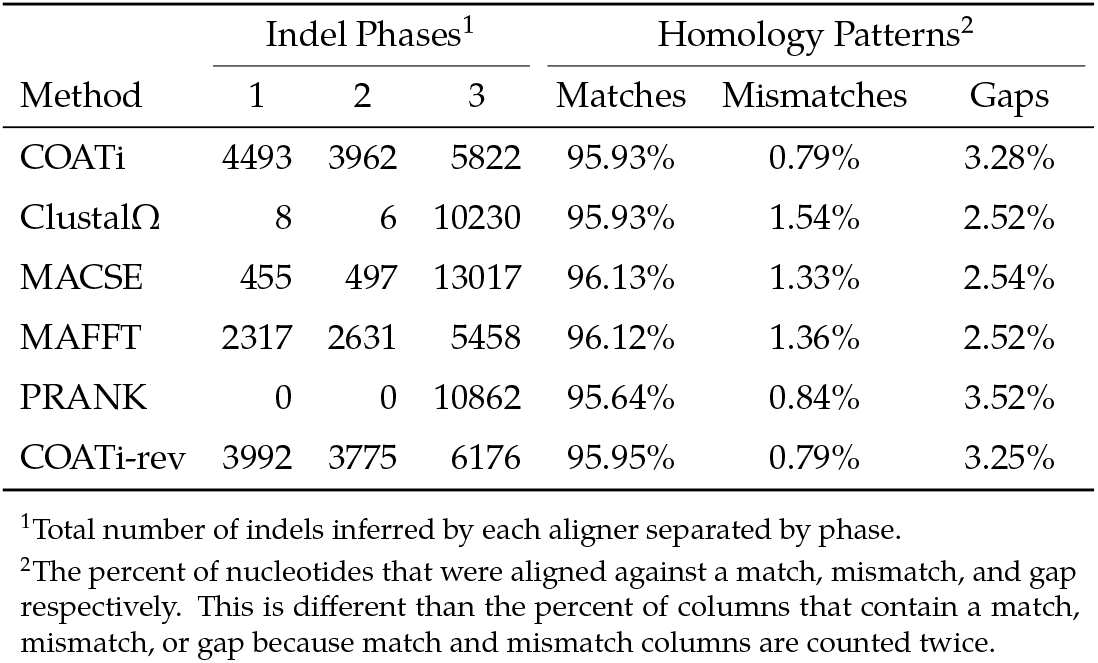
COATi produced more indels and fewer mismatches compared to other aligners.

### Evolutionary Distances

The distribution of evolutionary distances for COATi, COATi-rev, and PRANK were biologically reasonable for human-gorilla gene pairs, with means of 0.8%–0.9% (Fig. 4a). Conversely, the means for ClustalΩ, MACSE, and MAFFT were much higher, 1.5%–1.8%. These larger means are due to the fact that the right tails of the distributions for ClustalΩ, MACSE, and MAFFT were larger than the other methods. As expected, the alignment methods that “overaligned” sequences (produced more mismatches and fewer gaps) also produced higher evolutionary distances on average (Tab. 1 and Fig. 4a). The differences between the distances inferred by COATi and other methods also shows ClustalΩ, MACSE, and MAFFT producing higher distance estimates for a subset of sequences (Fig. 4b). COATi’s alignments produced significantly lower K2P distances than ClustalΩ, MACSE, MAFFT, and PRANK. Single-tailed Wilcoxon signed-rank tests on matched data produced p-values < 2.2 × 10^−16^. Additionally, COATi’s K2P distances did not depend on whether human or gorilla was used as the reference sequence (p-value of 0.4212 for a two-sided test). Additional analysis of the empirical alignments can be found in Supplementary Methods.

**Figure 4:**
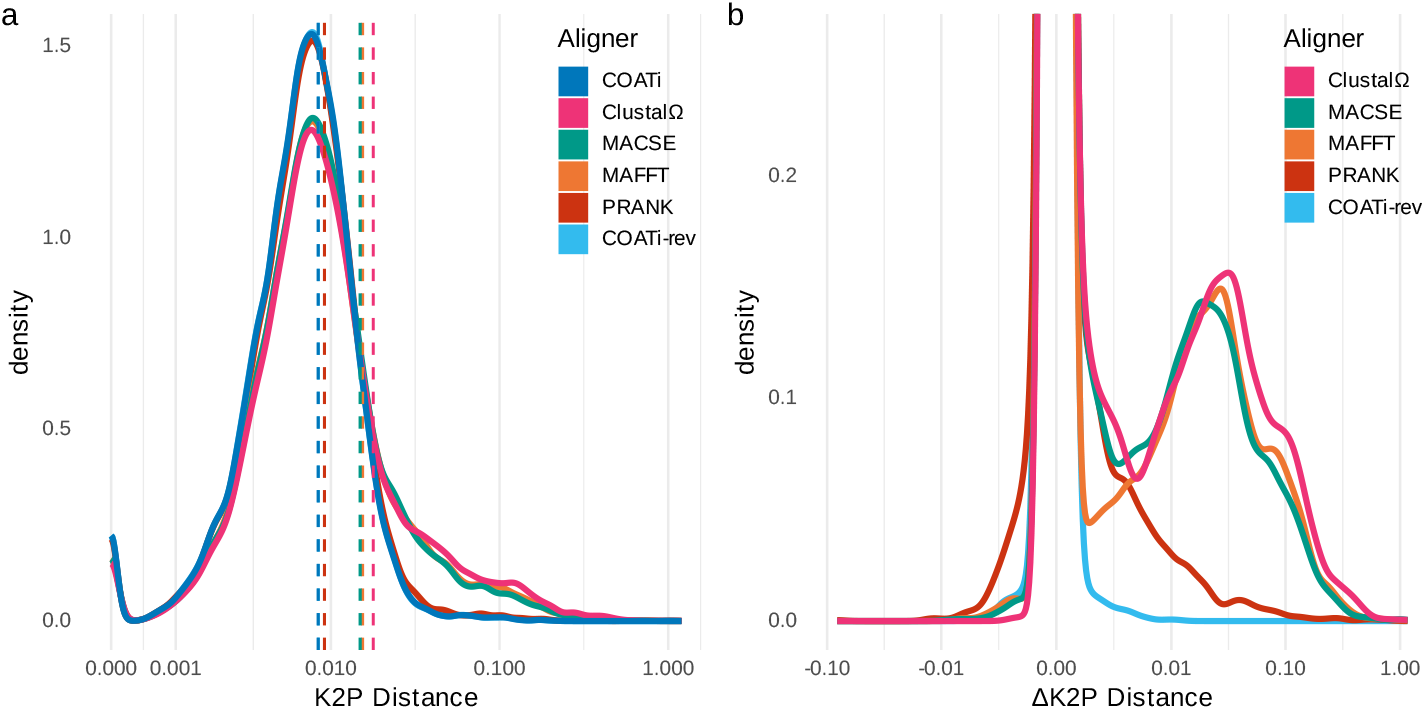
COATi’s alignments produce biologically reasonable evolutionary distances. (a) The distribution of K2P distances inferred from alignments generated by each method. The averages of each distribution are indicated by vertical lines. The averages are COATi = 0.0084, ClustalΩ = 0.0178, MACSE = 0.0149, MAFFT = 0.0154, PRANK = 0.0092, and COATi-rev = 0.0084. (b) The distribution of the differences between distances inferred by COATi and other methods. The X-axes of both plots have been pseudo-log transformed using the inverse hyperbolic sine.

### Selection

The vast majority of sequence pairs (12,344) were identified as showing evidence of negative selection when they were aligned using COATi, ClustalΩ, MACSE, MAFFT, PRANK, and COATi-rev (Fig. 5). Another 1087 sequence pairs were identified as showing evidence of positive selection when they were aligned using all 6 strategies. The most common pattern that showed disagreement between aligners were 91 sequence pairs that were identified as positively selected by COATi, PRANK, and COATi-rev. The most common singleton pattern was 64 sequence pairs identified as positively selected only by MAFFT. Notably, 36 sequence pairs were identified as positively selected and 9 sequence pairs were identified as negatively selected by by COATi and COATi-rev only. In total, COATi identified 1,367 (∼10%) sequence pairs as showing evidence of positive selection. ClustalΩ identified 1,228. MACSE identified 1,183. MAFFT identified 1,340. And PRANK identified 1,349. Note that we did not explore whether any of these inferences were biologically or statistically significant.

**Figure 5:**
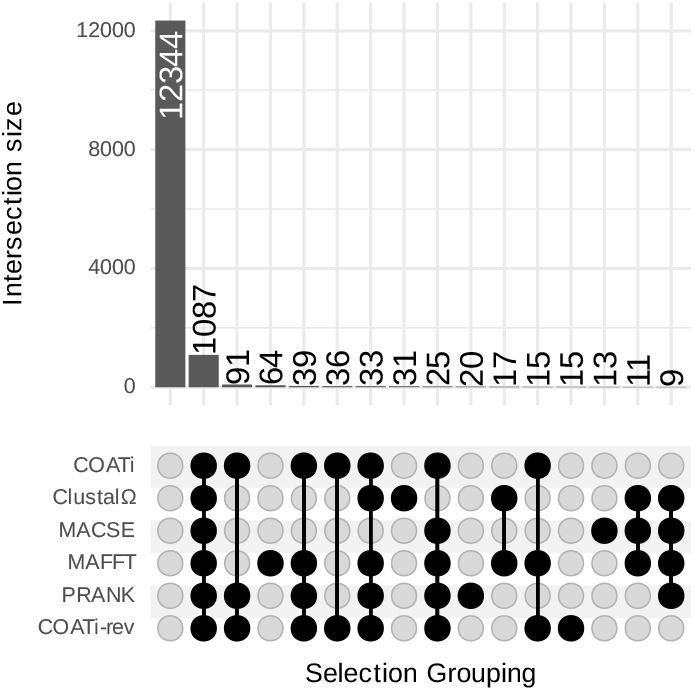
Aligners varied in which sequence pairs they identified as undergoing positive selection. In this UpSet plot, the bottom panel displays the 16 most frequent intersection patterns among aligners. A black circle represents positive selection. The most frequent pattern was that no aligner found positive selection while the second most frequent pattern was that all aligners found positive selection. Other patterns involved a disagreement between aligners about whether a sequence pair showed evidence of positive or negative selection. The top panel displays the number of sequence pairs in each grouping.

### Semi-Empirical Benchmark

COATi produced better alignments than ClustalΩ, MACSE, MAFFT, and PRANK when evaluated by our benchmark data set. It produced more best alignments, more perfect alignments, and less imperfect alignments (Tab. 2). COATi was significantly more accurate (lower *d*_*seq*_) at inferring the benchmark alignments compared to the other methods (Tab. 2). All p-values were significant according to the one-tailed, paired Wilcoxon signed-rank tests. (P-value of COATi vs PRANK was 5.1 × 10^−8^, and p-values of COATi vs the other aligners were < 2.2 × 10^−16^). Notably, the average alignment error of the second best protocol was four times larger than COATi’s. We also calculated the average distance of alignments between aligners and used principle coordinate analysis (PCoA) to project the resulting distance matrix into two dimensions (Fig. 6). COATi was the closest aligner to the benchmark, and other aligners tended to diverge from the benchmark in different directions. Because benchmark alignments were derived from gap patterns estimated by different aligners, we also performed separate PCoA analyses for each source of gap patterns (Supplementary Materials, Fig. S5). While all aligners, except ClustalΩ, do exceptionally well on their own patterns, COATi does exceptionally well on every set of patterns.

**Table 2:** COATi generates better alignments than other alignment algorithms on semi-empirical benchmarks.^1^

|  | COATi | ClustalΩ | MACSE | MAFFT | PRANK |
| --- | --- | --- | --- | --- | --- |
| Average Alignment Error ($d_{seq}$) | 0.23% | 2.65% | 1.37% | 1.67% | 1.02% |
| Number of Best<sup>2</sup> Alignments | 7050 | 3988 | 5226 | 5918 | 6128 |
| Number of Perfect<sup>2</sup> Alignments | 6795 | 3954 | 5138 | 5840 | 5860 |
| Number of Imperfect<sup>2</sup> Alignments | 850 | 3691 | 2507 | 1805 | 1785 |
| RMSE for K2P Distances | 0.000694 | 0.0920 | 0.0528 | 0.0561 | 0.0305 |
| Overestimated K2P Distances | 10.3% | 52.7% | 38.0% | 26.2% | 25.6% |
| F<sub>1</sub> Score for Negative Selection | 99.8% | 97.6% | 98.4% | 98.3% | 99.1% |
| F<sub>1</sub> Score for Positive Selection | 97.8% | 77.2% | 84.9% | 85.5% | 91.8% |

<sup>1</sup>Total number of sequence pairs in the benchmark was 8,261.
<sup>2</sup>Best alignments have the lowest $d_{seq}$ (including equivalent alignments). Perfect alignments have $d_{seq} = 0$ (including equivalent alignments). Imperfect alignments are alignments that are not perfect when at least one method found a perfect alignment.

**Figure 6:**
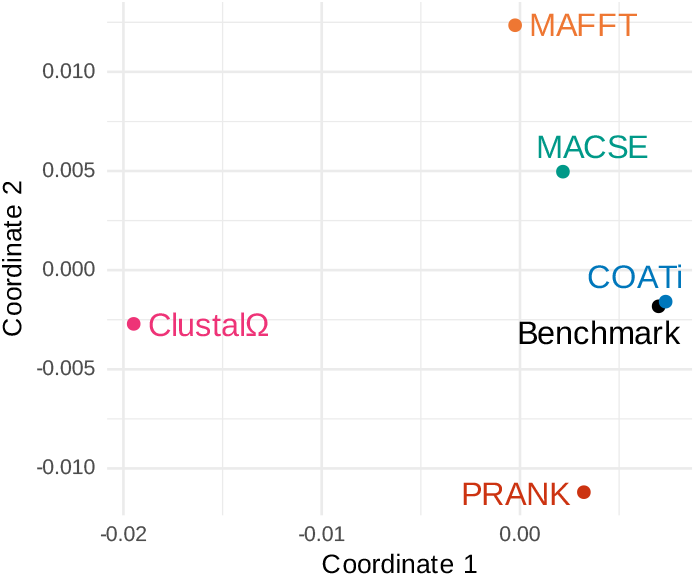
COATi’s alignments were closer to the semi-empirical benchmark than other methods. A principal coordinates analysis (PCoA) of the average alignment distance (*d*_*seq*_) between alignments generated by different methods.

For evolutionary distances, COATi had the lowest root-mean-square error (0.000694) relative to the benchmark alignments (Tab. 2). The second lowest aligner, PRANK, had a root-mean-square error that was over 400 times larger than COATi’s. COATi only overestimated 10% of its evolutionary distances, while other aligners overestimated 26%–53% of their evolutionary distances. Additionally, COATi more accurately inferred events of positive and negative selection (Tab. 2).

## Discussion

We have developed a statistical alignment program using finite-state transducers called COATi. On empirical data, COATi produced alignments that provided reasonable biological inferences. On semi-empirical, simulated benchmarks COATi was more accurate than the other aligners/alignment strategies that we tested: ClustalΩ’s amino-acid model, MACSE’s amino-acid aware nucleotide model, MAFFT’s DNA model, and PRANK’s codon model. Unlike standard codon or amino-acid alignment strategies, COATi supports aligning protein-coding sequences using all three phases of gaps. Consistent with Taylor et al. (2004) and Zhu (2022), our COATi results estimate that only 41% indels in protein-coding sequences between humans and gorillas are phase-3 indels. This indicates that alignment strategies that only support phase-3 gaps in protein-coding sequences are sub-optimally aligning the 59% of indels that are phase-1 or phase-2.

In addition to supporting all three phases of gaps, COATi’s other advantage over competing aligners in our tests is at least partially due to its default parameters fitting the human-gorilla coding sequences better than the other aligner’s default settings. Different data sets may show different results. However, because COATi uses a statistical model with biologically meaningful parameters, users can easily optimize COATi by using parameters estimated by previous studies or estimating parameters themselves (e.g. Zhu, 2022). This contrasts with other aligners that have limited ability to customize parameters and/or use parameters without biological interpretations.

COATi’s primary disadvantage is that its FST model was slower and consumed more resources than other methods. However, COATi’s marginal model produces results similar to the FST model, was the fastest method tested, and consumed reasonable amounts of memory (Supplementary Materials and García Mesa 2023). COATi is not a symmetric pairwise aligner, as reference sequences are more constrained than non-reference sequences. In order to test whether the choice of a reference sequence impact’s COATi’s results, we aligned our empirical sequence pairs with COATi using gorilla as the reference. We found no biologically or statistically significant differences between the results. The most noticeable difference was that, with gorilla as the reference, COATi rejected more sequence pairs due to the presence of ambiguous nucleotides in gorilla sequences.

After COATi, PRANK’s codon model was the second most accurate alignment method. Like COATi, PRANK also uses statistical models and supports aligning protein-coding sequences using codon models developed for phylogenetics. Both COATi and PRANK had reasonable distributions of evolutionary distances that lacked large right tails. This is likely because COATi and PRANK produced better alignments when tasked with aligning sequence pairs that contained non-orthologous exons. However, in contrast with COATi, PRANK’s codon model only permits phase-3 gaps and is very sensitive to frameshift artifacts. In fact, PRANK refuses to align any sequence using a codon model if its length is not a multiple of three.

ClustalΩ’s amino-acid model also only permits phase-3 gaps and is very sensitive to frameshift artifacts; however, it was the worst performing protocol that we tested. It had the highest average alignment error and also had difficulties correctly identifying positive selection and estimating evolutionary distances. — It even performed poorly on its own gap patterns. — ClustalΩ’s relatively poor performance likely derives from its default assumptions being a poor fit for our data set and also from the loss of information that occurred when we translated protein-coding DNA sequences into protein sequences.

We obtained better results when aligning in nucleotide space using both amino-acid aware MACSE and amino-acid agnostic MAFFT. These results indicate that aligning proteincoding sequences in nucleotide space works reasonably well if the sequences come from closely related species, such as humans and gorillas, as it supports all three gap phases and is robust to frameshift artifacts. If the sequences were further apart, we predict that amino-acid agnostic approaches would begin to produce unreasonable alignments.

We can look at distances between alignments generated by different aligners to understand the diversity of approaches use by our alignment protocols. Aside from the cluster of different COATi methods, we detected no other clusters in our analysis (Supplemental Materials Fig. S6). Different aligners tended to spread out from one another and in different directions from the benchmark. This indicates that each protocol that we utilized approached alignment inference from a different section of “aligner space”. While each aligner, except ClustalΩ, performed well on its own gap patterns, MACSE also did well on ClustalΩ and MAFFT gap patterns, and PRANK also did well on ClustalΩ patterns. COATi did well on all patterns.

COATi uses finite-state transducers to align pairs of sequences using a codon-aware statistical model. While COATi has multiple modes and models, here we have focused on evaluating the utility of COATi FST for estimating pairwise alignments. We have shown that COATi offers a biologically significant improvement over other methods when aligning pairs of sequences separated by short evolutionary distances in the presence of genomic artifacts. In other work, we have begun to explore application of COATi to more divergent sequences and the accuracy of the marginal approximation of the COATi FST (García Mesa, 2023).

COATi is under active development. We plan on extending it to support more complex gap models, e.g. mixtures of single-nucleotide and triple-nucleotide indel models and weighing gap openings to reflect known selection on indel phases (Zhu, 2022). We plan on improving its multiple-sequence alignment and alignment sampling capabilities, as well as implement new models for aligning long-read sequences of genes against reference genomes. Our goal is to develop COATi into a user-friendly suite of tools that will allow researchers to analyze more data with higher accuracy and facilitate the study of important biological processes that shape genomic data.

## Availability

The source code for COATi, along with documentation, is freely available on GitHub: https://github.com/CartwrightLab/coati and is implemented in C++. COATi is released under an MIT license. Additional information, code, and workflows to replicate the analysis can be found on GitHub: https://github.com/jgarciamesa/coati-testing and https://github.com/jgarciamesa/alignpair_letter.

## Acknowledgments

This research was funded by NSF award DBI-1929850. The authors would like to thank Profs. Marco Mangone, Ted Pavlic, Banu Oskan, Jay Taylor, and Jeremy Wideman for their helpful support on two separate PhD dissertation projects. The authors would also like to thank the associate editor and two reviewers for their helpful comments on earlier versions of this manuscript.

## Conflict of interest

none declared.

## Notes

https://github.com/CartwrightLab/coati

https://github.com/jgarciamesa/coati-testing

### Competing Interest Statement

The authors have declared no competing interest.

### Summary of Updates

We have broadened the scope of the manuscript from a software note to a larger research paper about our COATi software. Nearly all of the paper has been rewritten, expanded, and reconfigured to provide more details and background about our work. Importantly, we now use empirical results from the human-gorilla sequence pairs to initially compare aligners before we show results on semi-empirical benchmarks.

