## Supplementary Material for "COATi: statistical pairwise alignment of protein-coding sequences"

Supplementary Materials

Juan J. Garcia Mesa, Ziqi Zhu, Reed A. Cartwright

#### 1 Aligner Commands

We evaluated five different aligners. Below are the commands that we used to run them. We have abbreviated the commands for clarity, stripping out unimportant arguments. Complete workflows can be found in our coati-testing repository on Github.

- COATi: `coati alignpair -m tri-mg ...`; We used COATi two different ways. The primary way used human as the reference sequence. The secondary way, COATi-rev, used gorilla as the reference, via a wrapper script that reversed the order of sequences before alignment and reversed the order back afterwards.
- ClustalΩ v1.2.4: `clustalo --seqtype=Protein ...`; We wrapped ClustalΩ with a script that translates DNA sequences into amino acid sequences (including stops) before alignment. Any codons that were partial or containing ambiguous characters were translated as “X”. The script then aligned these translated sequences with ClustalΩ, and then created a DNA alignment that was consistent with the amino-acid alignment.
- MACSE v2.06: `java -jar macse.jar -prog alignSequences -seq human.fasta -seq_lr gorilla.fasta ...`; We wrapped MACSE with a script that created two temporary fasta files, one containing the human sequence and another containing the gorilla sequence. The human sequence was specified as the reliable sequence and the gorilla sequence was specified as the less-reliable sequence. MACSE uses “!” to mark gaps that result from frameshifts, and the wrapper script replaced these with “-”. Additionally, MACSE sometimes produced columns that only contained gaps, and these columns were removed.
- PRANK v.150803: `prank -codon ...`
- MAFFT v7.520: `mafft --preservecase --globalpair --maxiterate 1000 ...`

COATi can use different alignment models for pairwise alignment. Below are the commands that we used to run different models.

- tri-mg: `coati alignpair -m tri-mg ...`
- tri-ecm: `coati alignpair -m tri-ecm ...`
- mar-mg: `coati alignpair -m mar-mg ...`
- mar-ecm: `coati alignpair -m mar-ecm ...`

#### 2 FST Alignment Example

When using the triplet models (tri-mg and tri-ecm), COATi uses the OpenFST library to generate best alignments by composing the input and output sequences with the COATi FST model. While the COATi FST model is too large to display, we can show the result of a composition.

Fig. S1 shows a graph depicting the FST that results from composing the COATi FST with the input sequence "CTC" and the output sequence "CTG". Every path through this FST represents one possible way to align "CTC" and "CTG", and the sum of all weights along a path is the total weight of the respective alignment. Here, the weights of each arc are in negative-log space. Note that this graph has been optimized, and weight has been pushed towards the initial state. The weight of any specific arc may not be directly mapable to a weight described in the model.

Fig. S2 is the best alignment between "CTC" and "CTG", as determined by the shortest path algorithm. Figs. S1 and S2 were produced by the OpenFST library. A bold circle represents a starting node, and a double circle represents a termination node.

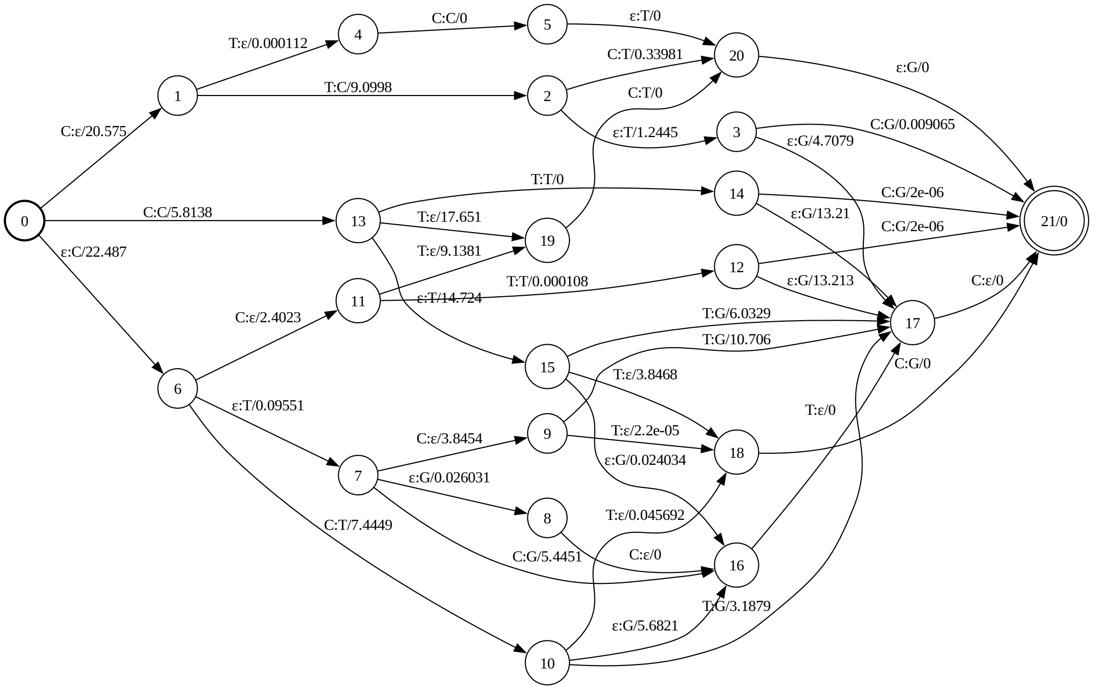

Figure S1: The FST of all possible alignments between "CTC" and "CTG".

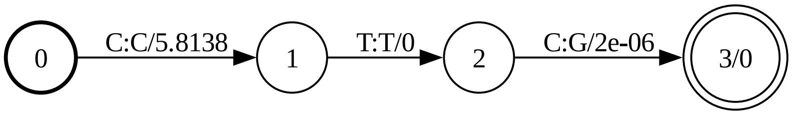

Figure S2: The best alignment of ”CTC” and ”CTG”.

### 3 Empirical Results

#### 3.1 Gap Patterns

##### 3.1.1 Lengths

We quantified the lengths of gaps produced by each method across the empirical dataset of human-gorilla protein-coding-sequence pairs (Tab. S1). Here the gap type is either "D" for gaps introduced into the gorilla sequence or "I" for gaps introduced in the human sequence. "COATi" refers to the full COATi FST model with a MG substitution model (i.e. tri-mg). "COATi-rev" refers to the same model, but using gorilla as the reference and human as the non-reference sequence. Gap lengths of 1–6 nucleotides were binned into their respective columns. Gap lengths longer than 6 nucleotides were binned into columns 7+, 8+, and 9+ depending on whether their lengths were 1, 2, or 3 nucleotides longer than a multiple of three.

We also quantified the lengths of gaps produced by different COATi models (Tab. S2).

Note that ClustalΩ gaps with lengths that are not a multiple of 3 are created by how our wrapper script handles DNA sequences with lengths that are not multiple of 3.

Table S1: Number of gaps introduced by each method separated by length and type.

| | | Gap Lengths | | | | | | | | |
|---|---|---|---|---|---|---|---|---|---|---|
| Method | Gap Type | 1 | 2 | 3 | 4 | 5 | 6 | 7+ | 8+ | 9+ |
| COATi | D | 106 | 99 | 651 | 94 | 94 | 226 | 1461 | 1386 | 3240 |
| COATi | I | 87 | 73 | 525 | 80 | 97 | 239 | 1399 | 1315 | 3105 |
| COATi-rev | D | 105 | 99 | 634 | 89 | 96 | 217 | 1425 | 1348 | 3169 |
| COATi-rev | I | 82 | 79 | 513 | 80 | 97 | 231 | 1360 | 1288 | 3031 |
| ClustalΩ | D | 1 | 0 | 887 | 1 | 1 | 400 | 3 | 7 | 3910 |
| ClustalΩ | I | 0 | 0 | 857 | 0 | 1 | 424 | 3 | 2 | 3747 |
| MACSE | D | 396 | 306 | 1237 | 5 | 3 | 657 | 71 | 54 | 4784 |
| MACSE | I | 60 | 27 | 1133 | 5 | 9 | 666 | 78 | 97 | 4381 |
| MAFFT | D | 204 | 156 | 714 | 147 | 108 | 276 | 544 | 509 | 2680 |
| MAFFT | I | 187 | 154 | 708 | 112 | 92 | 295 | 454 | 424 | 2642 |
| PRANK | D | 0 | 0 | 552 | 0 | 0 | 167 | 0 | 0 | 4812 |
| PRANK | I | 0 | 0 | 467 | 0 | 0 | 178 | 0 | 0 | 4686 |

Table S2: Number of gaps introduced by different COATi models separated by length and type.

| Model | Gap Type | Gap Lengths 1 | 2 | 3 | 4 | 5 | 6 | 7+ | 8+ | 9+ |
|---|---|---|---|---|---|---|---|---|---|---|
| TRI-MG | D | 106 | 99 | 651 | 94 | 94 | 226 | 1461 | 1386 | 3240 |
| TRI-MG | I | 87 | 73 | 525 | 80 | 97 | 239 | 1399 | 1315 | 3105 |
| TRI-ECM | D | 93 | 96 | 665 | 125 | 122 | 257 | 1438 | 1441 | 3203 |
| TRI-ECM | I | 86 | 103 | 540 | 105 | 122 | 254 | 1358 | 1370 | 3099 |
| MAR-MG | D | 102 | 106 | 646 | 95 | 89 | 224 | 1465 | 1399 | 3226 |
| MAR-MG | I | 84 | 83 | 526 | 80 | 101 | 238 | 1406 | 1313 | 3087 |
| MAR-ECM | D | 110 | 110 | 653 | 107 | 89 | 224 | 1424 | 1389 | 3250 |
| MAR-ECM | I | 86 | 89 | 523 | 77 | 99 | 249 | 1408 | 1310 | 3128 |
| DNA | D | 101 | 103 | 646 | 95 | 91 | 224 | 1446 | 1409 | 3206 |
| DNA | I | 79 | 79 | 520 | 78 | 99 | 239 | 1369 | 1316 | 3107 |

##### 3.1.2 Phases

We quantified the phases of gaps produced by different aligners and COATi models (Tab. S3 and S4). Phase 1 gaps begin after the 1st position in a codon in the reference sequence, phase 2 gaps begin after the 2nd position in a codon, and phase 3 begin after the 3rd position in a codon (i.e. between codons). Phase 3 gaps are also known as phase 0 gaps.

Note that ClustalΩ gaps with phases of 1 and 2 are created by how our wrapper script handles DNA sequences with lengths that are not multiple of 3.

Table S3: Number of gaps introduced by each alignment method separated by phase.

| Method | Gap Phases 1 | 2 | 3 |
|---|---|---|---|
| COATi | 4493 | 3962 | 5822 |
| COATi-rev | 3992 | 3775 | 6176 |
| ClustalΩ | 8 | 6 | 10230 |
| MACSE | 455 | 497 | 13017 |
| MAFFT | 2317 | 2631 | 5458 |
| PRANK | 0 | 0 | 10862 |

Table S4: Number of gaps introduced by different COATi models separated by phase.

| | Gap Phases | | |
|---|---|---|---|
| Model | 1 | 2 | 3 |
| TRI-MG | 4493 | 3962 | 5822 |
| TRI-ECM | 4160 | 4678 | 5639 |
| MAR-MG | 4130 | 3961 | 6179 |
| MAR-ECM | 4071 | 3863 | 6391 |
| DNA | 4228 | 3974 | 6005 |

#### 3.2 Homology Patterns

We quantified the homology patterns of residues for different aligners and COATi models (Tab. S5, Tab. S6, and Fig. S3). Here we define the match, mismatch, and gap percentages as the percent of nucleotides aligned against a match, mismatch, and gap respectively. Note that this is different than the percent of columns that contain a match, mismatch, or gap because match and mismatch columns are counted twice.

Table S5: Average homology percentages of alignments separated by alignment method.

| Method | Matches | Mismatches | Gaps |
|---|---|---|---|
| COATi | 95.93% | 0.79% | 3.28% |
| COATi-rev | 95.95% | 0.79% | 3.25% |
| ClustalΩ | 95.93% | 1.54% | 2.52% |
| MACSE | 96.13% | 1.33% | 2.54% |
| MAFFT | 96.12% | 1.36% | 2.52% |
| PRANK | 95.64% | 0.84% | 3.52% |

Table S6: Average homology percentages of alignments separated by COATi model.

| Model | Matches | Mismatches | Gaps |
|---|---|---|---|
| TRI-MG | 95.93% | 0.79% | 3.28% |
| TRI-ECM | 95.90% | 0.79% | 3.30% |
| MAR-MG | 95.93% | 0.79% | 3.28% |
| MAR-ECM | 95.94% | 0.80% | 3.26% |
| DNA | 95.93% | 0.79% | 3.27% |

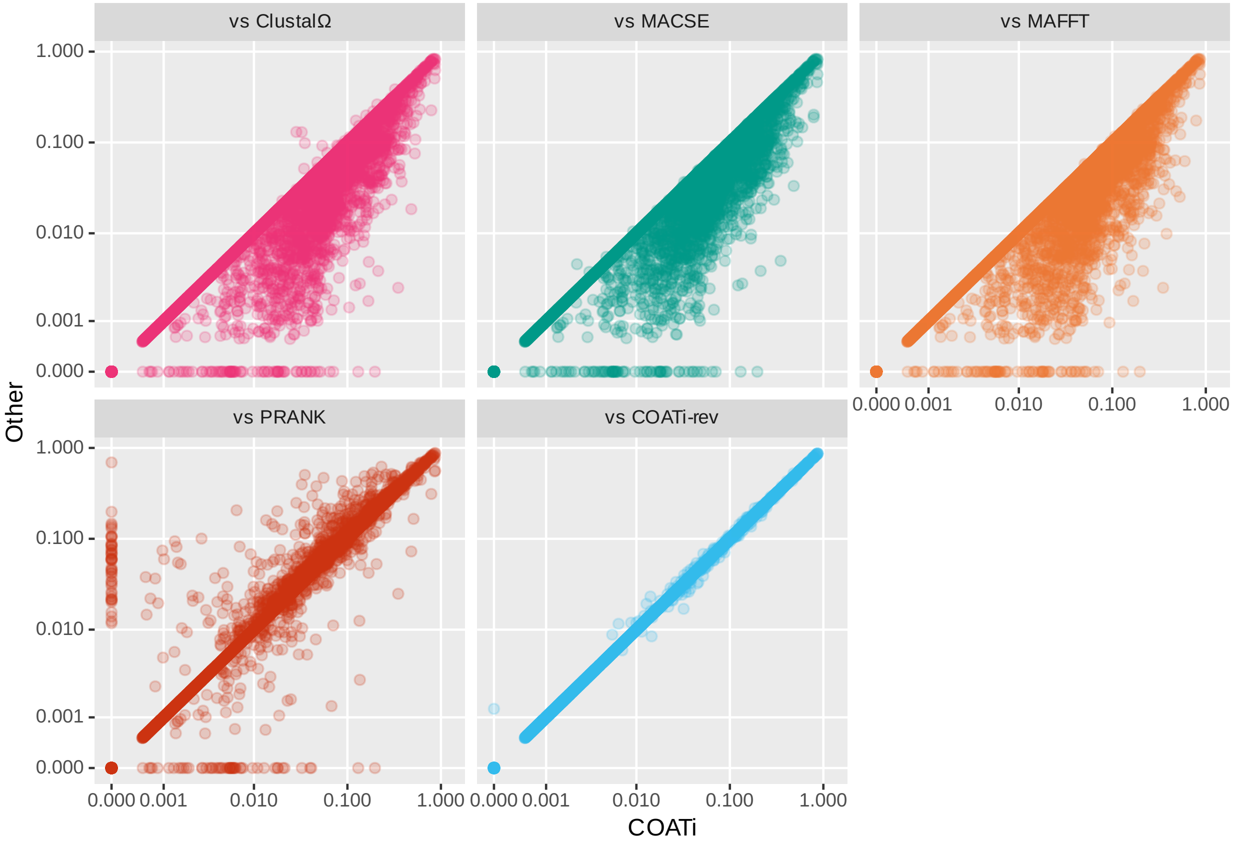

Figure S3: Gap fractions of COATi vs other aligners. Each panel is a scatter plot where the x coordinate is the gap fraction of a COATi alignment and y coordinate is the gap fraction of the corresponding alignment from another aligner.

#### 3.3 Evolutionary Distances

We quantified the evolutionary distances inferred by different aligners and COATi models (Fig. S4, Tab. S7, and Tab. S8).

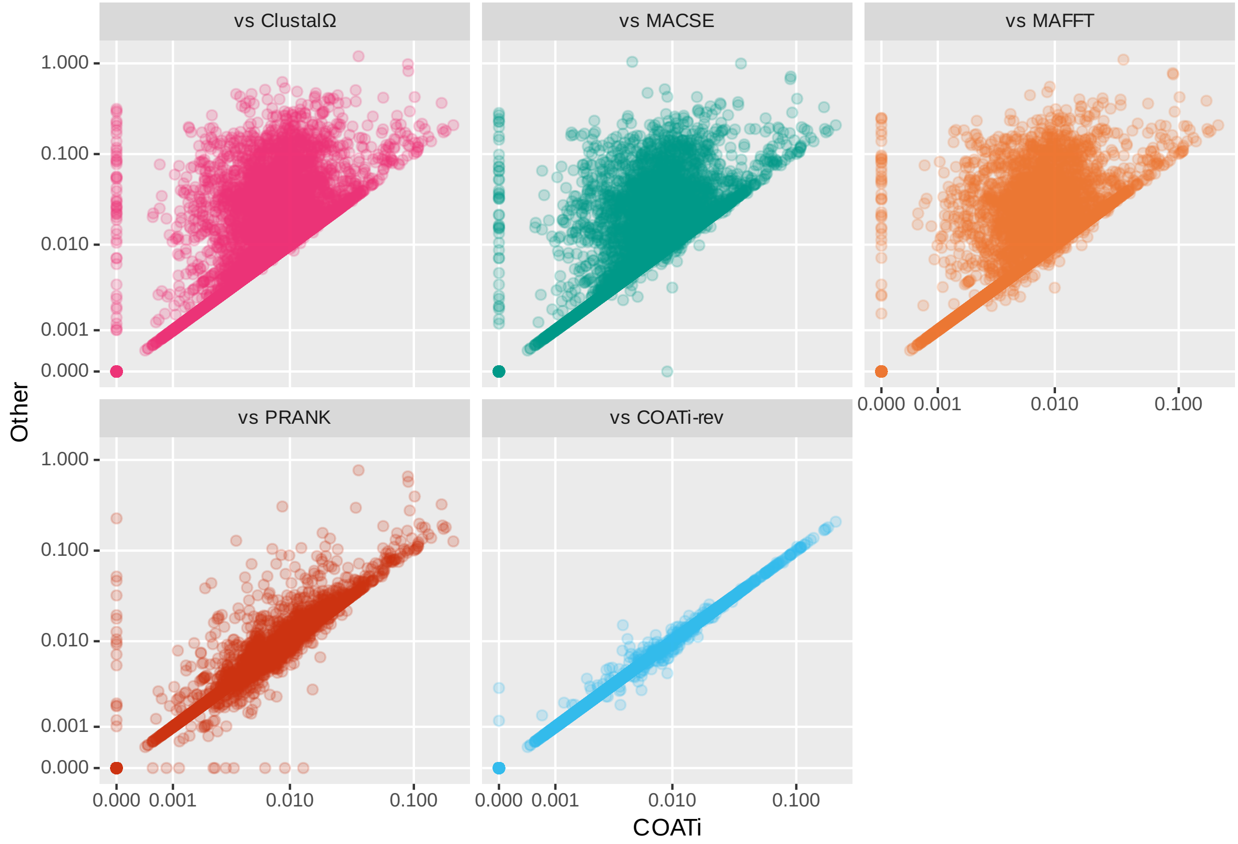

Figure S4: COATi produced shorter evolutionary distances than other aligners. Each panel is a scatter plot where the x coordinate is the K2P distance from a COATi alignment and y coordinate is the K2P distance from the corresponding alignment from another aligner.

Table S7: Average K2P distances of alignments for each method.

| COATi | COATi-rev | ClustalΩ | MACSE | MAFFT | PRANK |
|---|---|---|---|---|---|
| 0.0084 | 0.0084 | 0.0178 | 0.0149 | 0.0154 | 0.0092 |

Table S8: Average K2P distances of alignments for each COATi model.

| TRI-MG | TRI-ECM | MAR-MG | MAR-ECM | DNA |
|---|---|---|---|---|
| 0.0084 | 0.0084 | 0.0084 | 0.0084 | 0.0084 |

### 4 Benchmark Results

#### 4.1 Alignment Distances

For each sequence pair in the benchmark, we calculated the alignment distance ($d_{seq}$) between the benchmark alignment and the alignments generated by COATi, ClustalΩ, MACSE, MAFFT, and PRANK. We also calculated distances between all pairs of aligners. Our benchmark contained gap patterns extracted from alignments generated by different aligners. Figure S5 contains the results of a metric multidimensional scaling (principle coordinate analysis; PCoA) of the matrix of average distances between aligners, separated by what type of gap patterns was used in the benchmark alignment.

Figure S6 contains a principle coordinate analysis of each aligner, including different COATi models, across the entire benchmark dataset. COATi's different models produced similar alignments and cluster together along with the benchmarks.

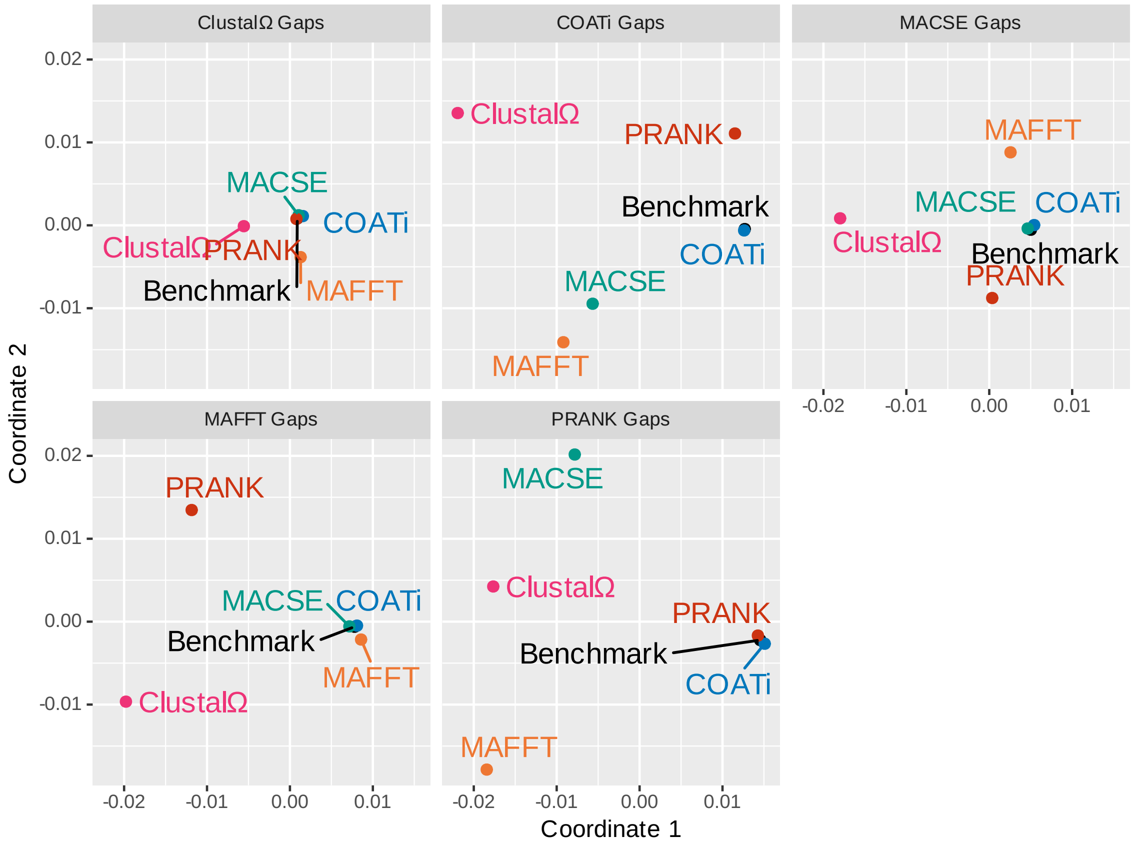

Figure S5: COATi produced accurate alignments regardless of whether the underlying gap pattern was extracted from an alignment generated by another program. Each panel is a metric multidimensional scaling of the average distances between aligners.

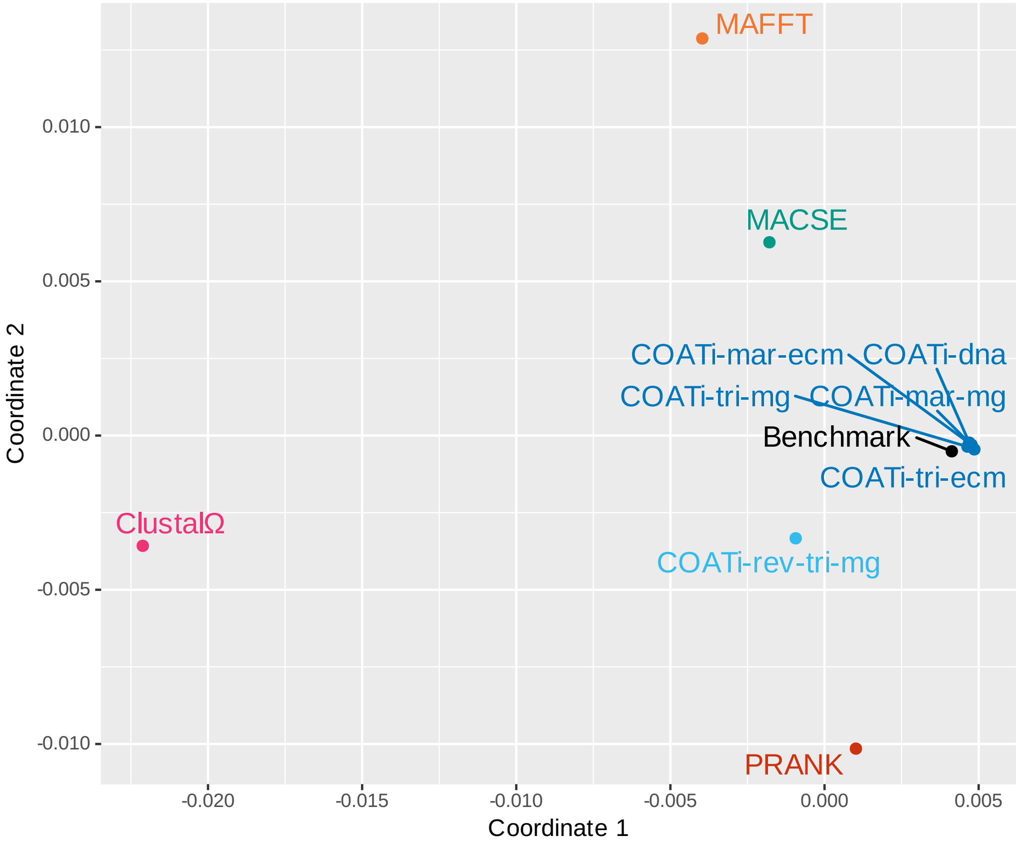

Figure S6: Different COATi models produce accurate alignments that are also similar to one another.
